# Arp2/3 and Unc45 maintain heterochromatin stability in Drosophila polytene chromosomes

**DOI:** 10.1101/679548

**Authors:** George Dialynas, Laetitia Delabaere, Irene Chiolo

## Abstract

Repairing DNA double-strand breaks (DSBs) is particularly challenging in pericentromeric heterochromatin, where the abundance of repeated sequences exacerbates the risk of ectopic recombination. In *Drosophila* Kc cells, accurate homologous recombination (HR) repair of heterochromatic DSBs relies on the relocalization of repair sites to the nuclear periphery before Rad51 recruitment and strand invasion. This movement is driven by Arp2/3-dependent nuclear actin filaments and myosins’ ability to walk along them. Conserved mechanisms enable the relocalization of heterochromatic DSBs in mouse cells, and their defects lead to massive ectopic recombination in heterochromatin and chromosome rearrangements. In *Drosophila* polytene chromosomes, extensive DNA movement is blocked by a stiff structure of chromosome bundles. Repair pathways in this context are poorly characterized, and whether heterochromatic DSBs relocalize in these cells is unknown. Here, we show that damage in heterochromatin results in relaxation of the heterochromatic chromocenter, consistent with a dynamic response in this structure. Arp2/3, the Arp2/3 activator Scar, and the myosin activator Unc45, are required for heterochromatin stability in polytene cells, suggesting that relocalization enables heterochromatin repair in this tissue. Together, these studies reveal critical roles for actin polymerization and myosin motors in heterochromatin repair and genome stability across different organisms and tissue types.

**Impact Statement:** Heterochromatin relies on dedicated pathways for ‘safe’ recombinational repair. In mouse and fly cultured cells, DNA repair requires the movement of repair sites away from the heterochromatin ‘domain’ *via* nuclear actin filaments and myosins. Here, we explore the importance of these pathways in *Drosophila* salivary gland cells, which feature a stiff bundle of endoreduplicated polytene chromosomes. Repair pathways in polytene chromosomes are largely obscure and how nuclear dynamics operate in this context is unknown. We show that heterochromatin relaxes in response to damage, and relocalization pathways are necessary for repair and stability of heterochromatic sequences. This deepens our understanding of repair mechanisms in polytenes, revealing unexpected dynamics. It also provides a first understanding of nuclear dynamics responding to replication damage or rDNA breaks, providing a new understanding of the importance of the nucleoskeleton in genome stability. We expect these discoveries to shed light on tumorigenic processes, including therapy-induced cancer relapses.

## Introduction

DNA is constantly under attack from endogenous and exogenous sources of damage, and DNA double-strand breaks (DSBs) are the most deleterious types of lesions because they interrupt the continuity of the DNA molecule. Repairing DSBs is particularly challenging in pericentromeric heterochromatin [1, 2] (hereafter ‘heterochromatin’), large blocks of highly repeated DNA sequences flanking the centromeres. Heterochromatin occupies nearly 30% of fly and human genomes [3-5], and is typically enriched for ‘silent’ chromatin marks (*e.g.,* H3K9me2,3 and its associated heterochromatin protein 1, or HP1). In *Drosophila melanogaster*, about half of these sequences are simple ‘satellite’ repeats, predominantly tandem 5-base pair sequences, repeated for hundreds of kilobases to megabases, while the rest are composed of scrambled clusters of transposable elements and about 250 isolated genes [3-5].

The two main pathways repairing DSBs are non-homologous end joining (NHEJ), which typically leaves small mutations at the cut site while rejoining the breaks [6], and HR, which repairs the lesion by copying information from a homologous template [7]. HR starts when DSBs are resected to form single-stranded DNA (ssDNA) filaments that invade ‘donor’ homologous sequences used as templates for DNA synthesis and repair [7]. In single copy sequences, a unique donor is present on the sister chromatid or the homologous chromosome, and repair is largely ‘error free’ [7]. In heterochromatin, thousands to millions of potential donor sequences associated with distinct chromosomes can initiate inter-chromosomal recombination or unequal sister chromatid exchange, resulting in translocations, deletions, duplications, and formation of dicentric or acentric chromosomes [1, 8-11]. Despite this danger, HR is a primary pathway for heterochromatin repair in both *Drosophila* and mammalian cells [10-16], and specialized mechanisms enable ‘safe’ repair in heterochromatin while preventing aberrant (ectopic) recombination.

In *Drosophila* cultured cells, where heterochromatin forms a distinct nuclear ‘domain’ [13, 17, 18], HR starts inside the domain with resection [10, 13, 14, 19] (Figure 1), while subsequent repair steps are temporarily halted [10, 13, 20]. This block to HR progression relies on Smc5/6, which is recruited to heterochromatin by HP1a [10, 13, 20], and three SUMO E3 ligases: dPIAS and the Smc5/6 subunits Nse2/Qjt and Nse2/Cerv [10, 13, 20]. Next, the heterochromatin domain expands while repair sites display a striking relocalization to the nuclear periphery, where repair progresses [10, 13, 14, 19]. Relocalization relies on a remarkable network of nuclear actin filaments assembled at repair sites by the actin nucleator Arp2/3, and its activators Scar and Wash [11]. Filaments extend toward the nuclear periphery [11], consistent with a role of filaments as ‘highways’ for relocalization. Relocalization also requires three nuclear myosins: Myosin 1A, Myosin 1B, and Myosin V, and myosin’s ability to ‘walk’ along actin filaments [11], revealing transport mechanisms similar to those acting in the cytoplasm [21-23]. In agreement, relocalization of heterochromatic DSBs to the nuclear periphery is characterized by directed motions [11, 24]. Arp2/3 and myosin recruitment to repair sites requires the early DSB signaling and processing factor Mre11, and the heterochromatin protein HP1a [11], suggesting the combination of these components as a mechanism for targeting the relocalization machinery specifically to heterochromatic DSBs. Additionally, Smc5/6 recruits Unc45 myosin activator to heterochromatic repair sites [11], suggesting this step as a critical switch to promote myosin ‘walk’ along the filaments and DSB relocalization [25].

**Fig. 1:**
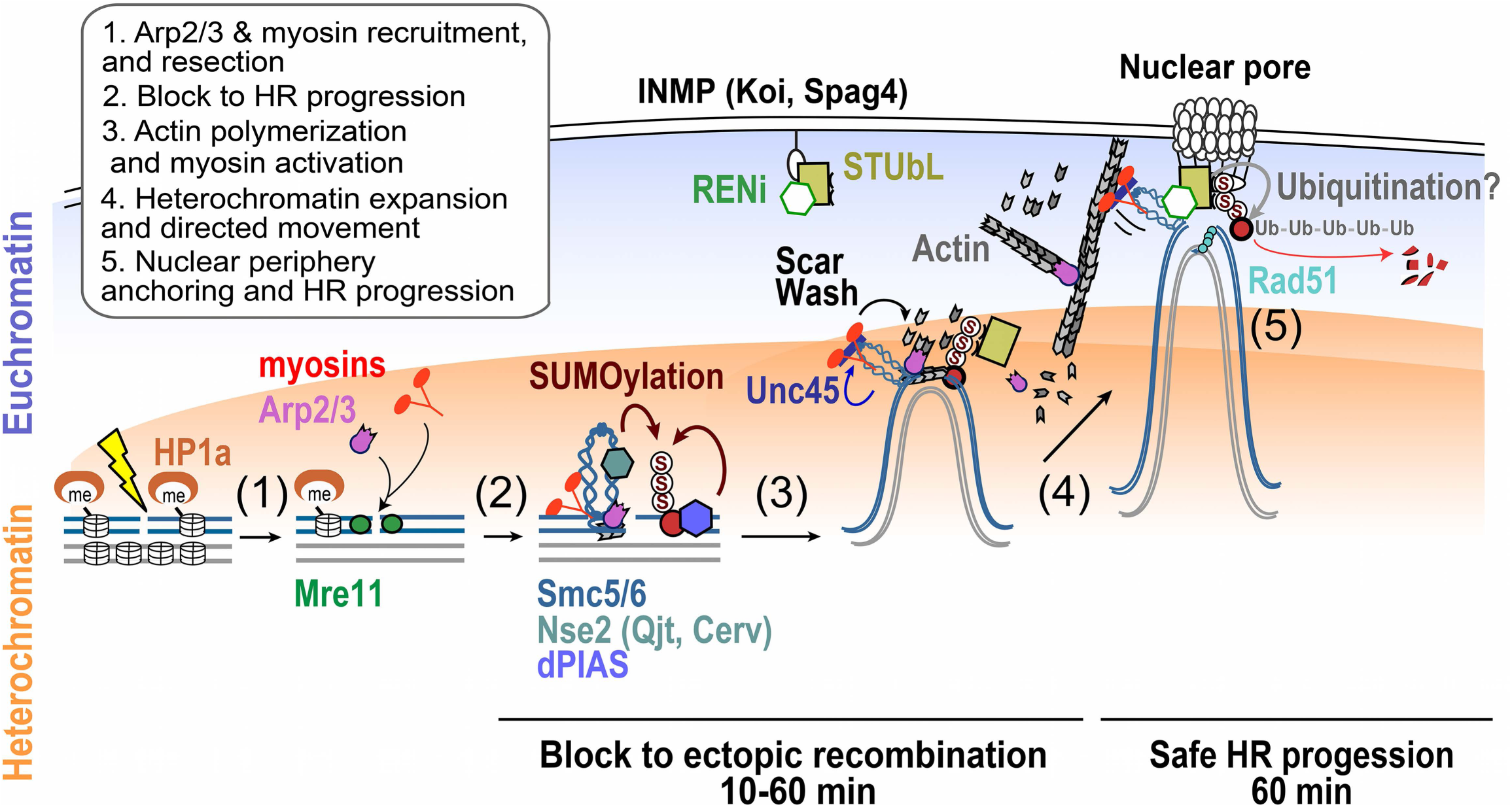
Schematic view of the molecular mechanisms required for heterochromatin repair in *Drosophila* cultured cells. See Introduction for details; the main repair steps are numbered and indicated.

Inactivating the relocalization pathway results in aberrant recombination and widespread genomic instability [10, 11, 13, 20], revealing its importance to genome integrity. Relocalization likely promotes ‘safe’ HR repair while preventing aberrant recombination, by isolating DSBs and their repair templates (on the homologous chromosome or the sister chromatid) away from non-allelic (ectopic) sequences before strand invasion [10, 11, 13, 20] (reviewed in [2, 26, 27]). Notably, *Drosophila* homologous chromosomes are paired in interphase, thus both sister chromatids and homologous chromosomes are readily available for DSB repair. In agreement, disruption of sister chromatid cohesion or homologous pairing results in heterochromatin repair defects [11].

Relocalization of heterochromatic DSBs and expansion of the domain also occurs in mouse cells [2, 11, 16, 28, 29], suggesting conserved pathways [26, 27]. However, in mouse cells, heterochromatic DSBs do not appear to reach the nuclear periphery and repair sites are detected at the periphery of DAPI-bright regions, where Rad51 is recruited [2, 11, 16, 29]. Nevertheless, Arp2/3, actin polymerization and myosins also mediate relocalization and repair of heterochromatic DSBs and genome stability in this system [11]. Additionally, data in *Drosophila* and human cells point to a role for Arp2/3 in DSB clustering and HR repair in euchromatin [11, 25], also suggesting the importance of actin-driven relocalization pathways in local dynamics.

In flies, a distinct nuclear organization characterizes polytene chromosomes, which contain hundreds to thousands of copies of chromosomes associated lengthwise to form a cable-like structure (reviewed in [30, 31]). Polytene chromosomes are found in several tissues of diptera, and originate from successive replications of each chromatid without segregation (endocycles). In larval salivary glands of *Drosophila,* cells undergo approximately 10 endoreplication cycles, resulting in 1024 copies of most loci [30]. Given the large size, polytene chromosomes have been an invaluable tool for mapping gene position, chromosome rearrangements, and chromatin states (*i.e.* active *vs* inactive regions) across the genome [31]. In these cells, DNA staining shows a typical banding pattern, with active genes localized in interbands, and silent, more compact, chromatin in darker bands [31-36].

Late-replicating regions (*i.e.,* pericentromeric heterochromatin, and silenced genes along chromosome arms called intercalary heterochromatin [32, 34, 37-39]) remain largely underreplicated in polytene chromosomes. This results in stalling and collapse of forks entering the domain, followed by mutagenic repair [40-42]. In agreement, whole genome sequencing revealed large deletions at intercalary heterochromatin [43]. Similarly, cytological preparations reveal filamentous structures connecting under-replicated regions, corresponding to ectopic exchanges [40, 43, 44]. In pericentromeric heterochromatin, loss of the silencing component Su(var)3-9 results in accumulation of DNA damage [9] and formation of extra-chromosomal circles (ECCs) of satellite sequences in salivary gland cells [8], also reflecting mutagenic repair. The extent of pericentromeric heterochromatin repair in polytene chromosomes of wild-type flies is unclear, and a direct investigation of DSB repair in these regions is lacking.

At the cytological level, pericentromeric heterochromatin of polytene chromosomes clusters in a large domain called ‘chromocenter’ from which chromosome arms depart. Notably, mobilization of repair sites is central for heterochromatin repair in mouse and *Drosophila* cultured cells, but the ‘rigid’ structure of polytene chromosomes is potentially a major impediment to chromatin dynamics. Here, we investigate the relevance of the relocalization pathway in polytene cells. We show that chromocenters expand in response to ionizing radiation (IR), revealing a dynamic reorganization of the domain during repair. Intriguingly, removing relocalization pathway components Arp2/3, Scar, or Unc45, results in heterochromatin and rDNA instability in polytene chromosomes. These data support a role for relocalization in heterochromatic DSB repair in salivary gland cells, and identify a new pathway for repair and stability of heterochromatin in polytene chromosomes.

## Materials and methods

### Fly stocks and crosses

*Drosophila* stocks were maintained on standard media at 25°C, prepared as described in [45]. Stocks were obtained from BDSC (http://fly.bio.indiana.edu) or VDRC (www.vdrc.at) and are: Arp3 (BDSC #32921) y[1] sc[*] v[1]; P{y[+t7.7] v[+t1.8]=TRiP.HMS00711}attP2; Scar, (BDSC #31126) y[1]v[1];P{y[+t7.7] v[+t1.8]=TRiP.JF01599}attP2; Act5c-GAL4 (BDSC #4414) y[1] w[*]; P{w[+mC]=Act5C-GAL4}25FO1/CyO, y[+]; Unc45 (VDRC #v108868) P{KK101311}VIE-260B. *su(var)3-9^null^* trans-heterozygous mutant was *su(var)3-9^6/17^*previously described in [8]. *yry* (kind gift from G. Karpen) was used as a control in Supplementary Fig. 2. To obtain 3^rd^ instar larvae for salivary gland dissection, RNAi lines or a control *w*^*1118*^ were crossed to the Act5c-GAL4 line (rebalanced with CyO-GFP) and non-GFP larvae were picked for dissection [46]. Efficiency of RNAi depletions were previously validated [11]. mGFP-Mu2 flies (kind gift from J. Mason) were previously described [47].

**Fig. 2:**
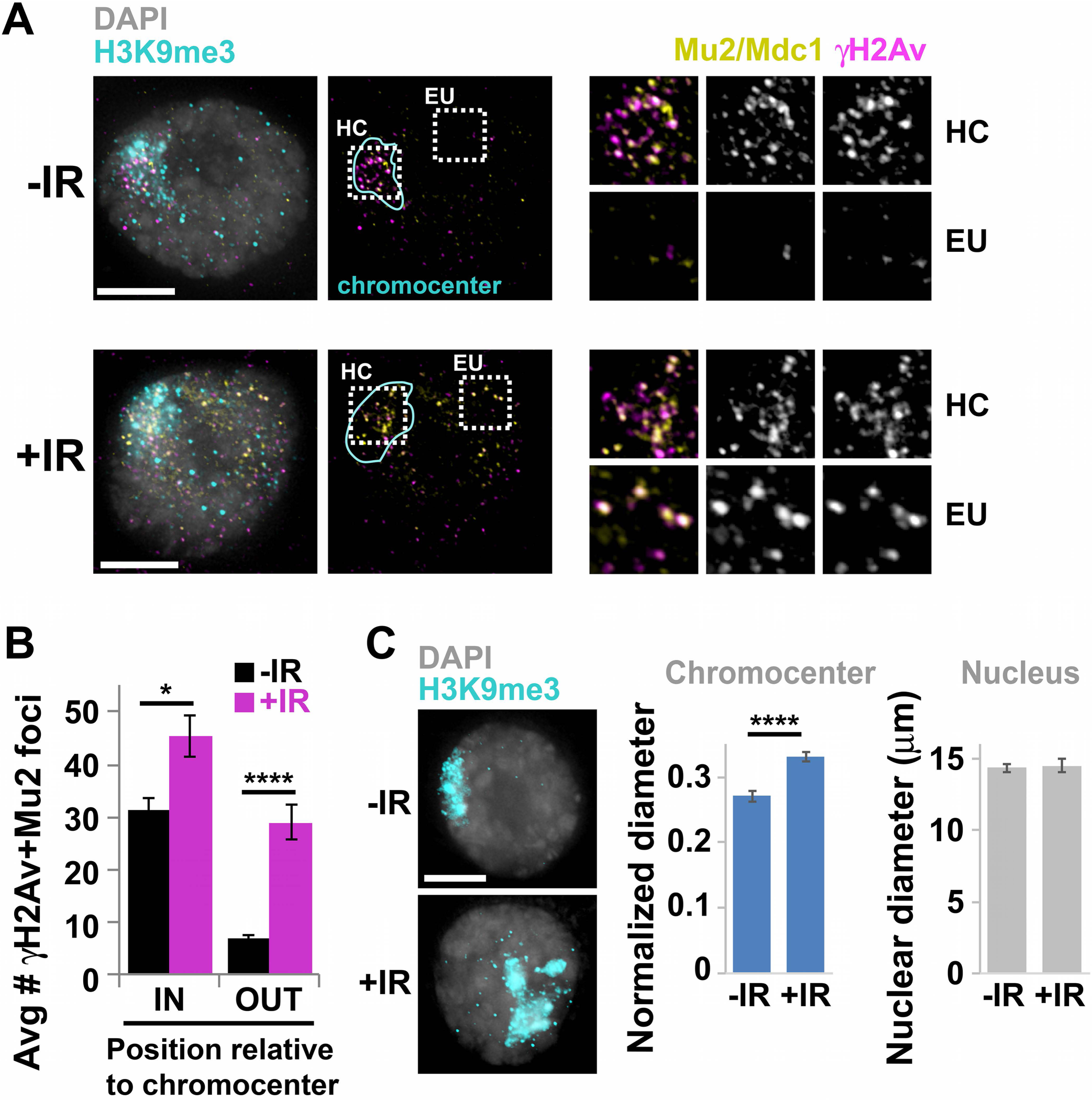
A) IF analysis and B) quantification of repair foci containing both γH2Av and Mu2/Mdc1 signals inside (HC, heterochromatin) and outside (EU, euchromatin) the chromocenter before (-) and after (+) treatment with 20 Gy IR. *P=0.01; \*\*\*\**P*<0.0001 relative to untreated (-IR); two-tailed Mann-Whitney test; n=17 salivary gland cells from 2 independent experiments. The average nuclear size for the cells quantified here is shown in Supplementary Figure 1. C) IF analysis and quantification of the average size of the chromocenter (H3K9me3-positive domain) and of the nucleus of salivary gland cells before (-) and after (+) treatment with IR. \*\*\*\**P*<0.0001 relative to untreated (-IR), unpaired t test with Welch’s correction, n>35 salivary gland cells. Error bars: SEM. Scale bars = 5µm.

### IF, FISH, imaging, quantifications

Salivary glands were dissected from 3rd instar larvae in PBS with Triton 0.15% (PBST) and fixed in PBST plus 4% PFA for 15 min, before IF. FISH was performed on whole mount tissues as described in [8], and probe hybridization was done in microtubes. FISH (including probe design) was done as described in [48, 49]. IF, imaging and image processing for fixed tissues was done as described in [13]. The number of pH2Av+Mu2/M2c1 signals was calculated in volume reconstructions of salivary gland cells, relative to the chromocenter (H3K9me3-positive signal). For the calculation of nuclear and chromocenter diameters, several Z-stacks encompassing salivary glands were used for volume reconstructions using SoftworX. Nucleus and chromocenter diameter were calculated along the XY plane with the ‘measure distance’ function in SoftworX.

### Antibodies

Primary antibodies used in *Drosophila* cells were: anti-γH2Av (1:1000, Rockland, 600-401-914); anti-GFP (1:1000 Aves Lab, GFP-1020); anti-H3K9me2 (1:750, Wako Chemicals, MABI0307, 302-32369). Secondary antibodies for IF were from Life Technologies and Jackson Immunoresearch. Antibodies were previously validated [10, 13, 50].

### Statistical analyses

All statistical analyses were performed using Prism6 (Graphpad Software).

## Results

Much progress in our understanding of heterochromatin repair comes from studying IR-induced damage in *Drosophila* and mouse cells. The recruitment of repair components to DSBs results in cytologically visible foci [51], and the distribution of those relative to the heterochromatin domain enables the spatial and temporal characterization of repair responses [2, 10, 11, 20, 52, 53]. However, polytene cells are characterized by a large number of breaks from replication fork collapse that might hinder detection of additional damage from IR. We tested this directly in flies expressing a GFP-tagged version of Mu2/Mdc1, an early component of the DSB response. Tissues were fixed before and 15 min after 20 Gy IR, and DSBs were detected using antibodies for GFP and the phosphorylated form of H2Av (γH2Av), a histone mark deposited in response to DNA breaks and recognized by Mu2/Mdc1. Despite some background signal in these large nuclei, repair sites are easily detected as foci containing both Mu2 and γH2Av signals (Figure 2A). Before IR, cells contain few repair foci outside the chromocenter, and a large number of repair foci associated with the chromocenters (Figure 2A). IR treatment induces a major increase of repair foci outside the chromocenters, but only a modest increase inside (Figure 1 A,B). Higher IR doses result in excessive level of foci that cannot be reliably counted (not shown). Intriguingly, IR also results in a significant volume increase of the chromocenter (Fig. 1C), resembling heterochromatin domain expansion in cultured cells [13]. This is consistent with a global relaxation of heterochromatin in response to DNA damage.

Given that the high level of pre-existing damage in heterochromatin limits the detection of IR-induced events, we focused on damage responses to endogenous heterochromatic breaks derived from endocycles. We used a genetic approach to investigate the relevance of relocalization pathway components to heterochromatin stability in salivary gland cells. Previous studies showed that loss of the histone methyltransferase Su(var)3-9, responsible for H3K9 di- and tri-methylation in heterochromatin, results in rDNA instability detected as multiple nucleoli in salivary gland cells [8]. Notably, nucleoli are enriched for repeated DNA sequences, and are at least partially silenced, retaining some features of pericentromeric heterochromatin [8]. Similarly, we detected additional heterochromatin signals in salivary gland cells of *su(var)3-9*^*null*^ mutants (Supplementary Figure 2). Multiple nucleoli and heterochromatin domains likely result from repair defects in *su(var)3-9* mutants [8, 9, 13], leading to sequence amplification and formation of extrachromosomal circles (ECCs) of repeated DNA sequences [8]. These phenotypes are particularly evident in salivary gland cells given that those do not segregate the genetic material, thus retaining extra-chromosomal DNA copies in the nucleoplasm [8].

Remarkably, RNAi depletion of relocalization pathway components Arp3, Scar, and Unc45 results in multiple nucleoli and additional heterochromatin signals in salivary gland cells (Figure 2), compared to control RNAi flies. This is consistent with amplification and excision of pericentromeric and rDNA sequences in the absence of relocalization pathway components.

We directly tested the effect of these depletions on heterochromatin stability by investigating the number and nuclear position of satellite signals, using fluorescence in situ hybridization (FISH). Previous studies showed that *su(var)3-9* mutation results in additional copies of the large 359bp satellite associated with the X chromosome, along with longer distance between FISH signals [8]. These reflect the formation of ECCs of satellite DNA [8]. We confirmed this observation, and extended it to AACAC and AATAT satellites (Figure 3). Similarly, RNAi depletion of Arp3, Scar and Unc45 results in higher numbers of satellite signals and longer distance between the signals, relative to control RNAi (Figure 3). We conclude that loss of relocalization pathway components results in additional copies of satellite sequences, including amplification and excision of satellite DNAs. Importantly, the nuclear size of cells quantified in these experiments is consistent across different RNAi depleted and mutant flies, and in IR-treated relative to untreated tissues (Figure 2,3; Supplementary Figure 1, 2), ruling out that the observed effects result from differences in the number of endocycles.

**Fig. 3:**
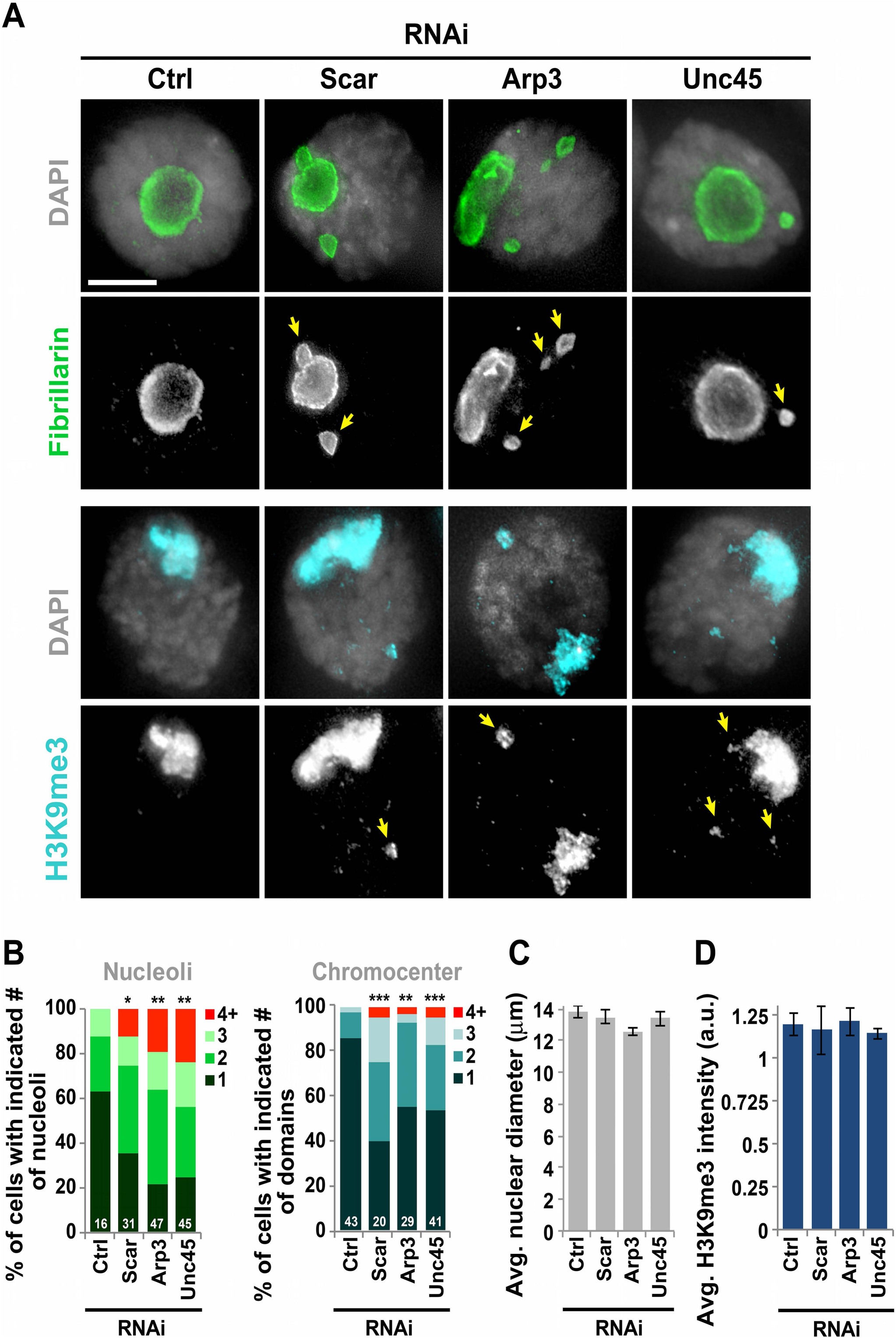
A) IF analysis and B) quantification of the number of nucleoli (fibrillarin-positive signals [8], arrows) and heterochromatin domains (H3K9me3-positive signals, arrows) in salivary gland cells of control (Ctrl, Act5C-Gal4/CyO) or RNAi depleted flies, as indicated. In B), numbers next to each graph indicate the number of ‘domains’ quantified; numbers inside the graph are the number of cells quantified. *P=0.038, one-tailed Mann-Whitney test; \*\**P*<0.045, \*\*\**P*<0.0008, two-tailed Mann-Whitney test. C) Quantification of the nuclear diameter for the cells shown in B,D) Quantification of the average intensity of H3K9me3 signals in the chromocenters of cells shown in B. a.u.: arbitrary units. Error bars: SEM. Scale bars = 5µm.

Additionally, Su(var)3-9 mutations result in a significant reduction of H3K9me3 signals in heterochromatin [8, 13], reflecting the importance of Su(var)3-9 in H3K9me3 establishment and maintenance. Conversely, RNAi depletion of Arp3, Scar, or Unc45 did not affect H3K9me3 intensity (Figure 3D), consistent with a role of relocalization components in heterochromatin maintenance downstream from Su(var)3-9. Together, these studies reveal the importance of relocalization pathway components in heterochromatin repair and stability in polytene chromosomes.

## Discussion

In *Drosophila* cultured cells, ‘safe’ HR repair of heterochromatic DSBs relies on the relocalization of repair sites to the nuclear periphery, and this requires the formation of Arp2/3-dependent nuclear actin filaments and the ability of myosins to ‘walk’ along the filaments [10, 11, 13]. Here we show that the actin polymerizing complex Arp2/3, its activator Scar, and the myosin chaperone Unc45 are also required for heterochromatin repair in polytene chromosomes. This is surprising, given that polytene chromosomes form rigid structures that likely interfere with chromatin dynamics. However, substantial opening of the chromatin is observed in highly transcribed genes (i.e., DNA ‘puffs’ [54, 55]), thus local dynamics are not uncommon in these tissues. We also detected a significant increase of chromocenter volume in response to DNA damage, similar to the expansion of the heterochromatin domain observed in *Drosophila* and mouse cultured cells [13]. We suggest a model where heterochromatin relaxation facilitates Arp2/3 and myosin-driven relocalization of repair sites to outside the chromocenter of polytene chromosomes to minimize the risk of ectopic recombination during DSB repair (Figure 4).

**Fig. 4:**
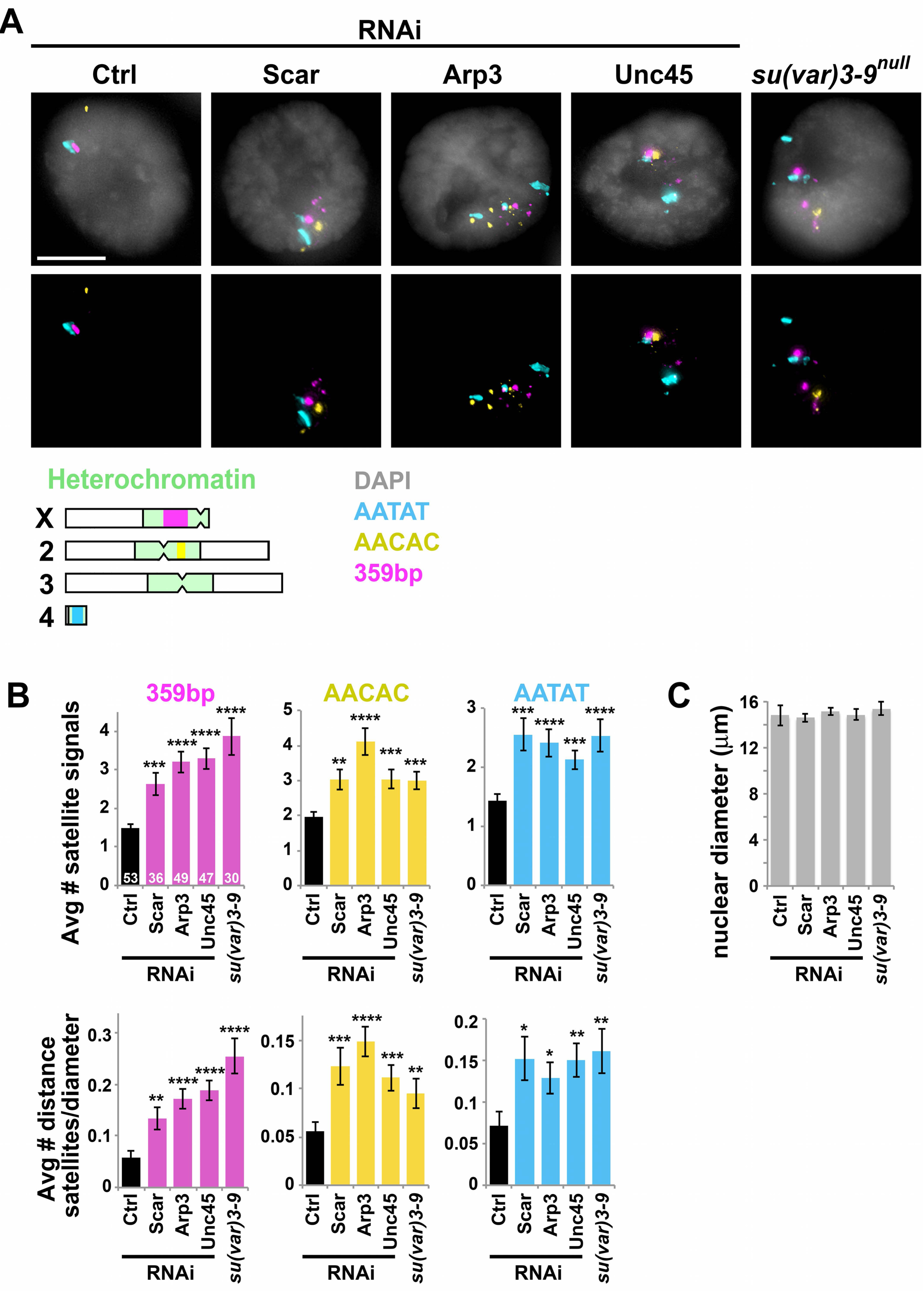
A) FISH analysis (top), schematic representation of the position of the corresponding satellites (bottom), and B) quantification of indicated satellite signals in salivary glands of control flies (Ctrl, Act5C-Gal4/+), flies undergoing indicated RNAi depletions, or *su(var)3-9*^*null*^ mutants. *P<0.03, **P<0.006, ***P<0.001, ****P<0.0001 relative to Ctrl; two-tailed Mann-Whitney test. Error bars: SEM. The number of cells quantified is indicated in the graph. C) Quantification of the nuclear diameter for the cells analyzed in B).

**Fig. 5:**
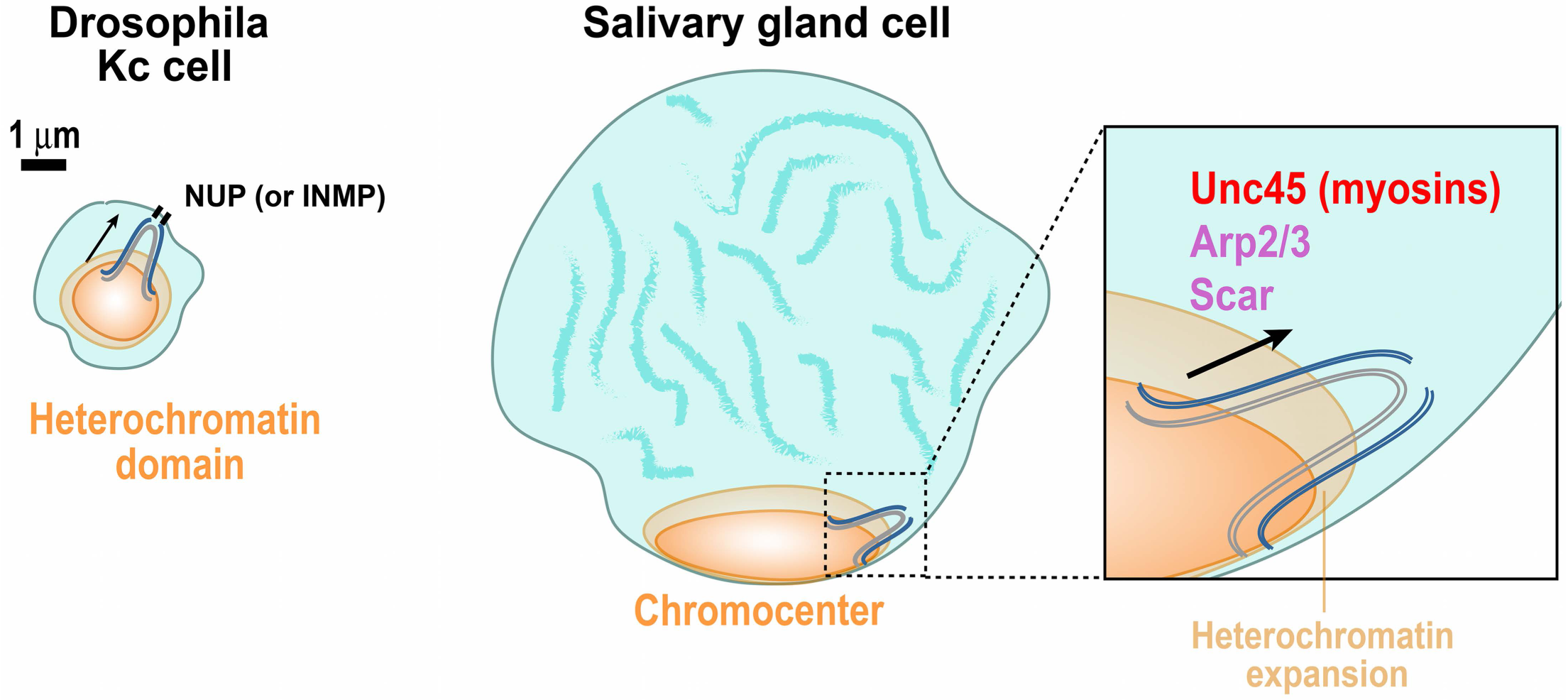
Model for the relocalization of heterochromatic DSBs in *Drosophila* cultured cells and salivary gland nuclei.

Whether heterochromatic repair sites reach the nuclear periphery in these large nuclei remains unclear. DSBs are frequently found at the periphery of the chromocenter in *Drosophila* polytene cells ([42] and data not shown), supporting a ‘local relocalization’ model in these cells. Thus, DSBs could relocalize to the chromocenter periphery rather than to the nuclear periphery, similar to what is observed in mouse cells. However, chromocenters are also typically associated with the nuclear periphery [56, 57], minimizing the distance repair sites would need to travel to reach this location; more studies are needed to determine the nuclear position where repair continues in this tissue, the pathway responsible for repair restart, and repair outcomes.

Intriguingly, nuclear actin polymers have been previously detected in *Drosophila* nurse cells, which undergo polytenization similar to salivary gland cells [58-60]. Nurse cells also accumulate DNA damage during endocycles and heterochromatin replication [61], suggesting a role for nuclear actin structures in DNA repair of polytene chromosomes in nurse cells.

Our data also revealed that Arp2/3, Scar and Unc45 are required for nucleolar stability, suggesting a role for these components in rDNA repair. Damaged rDNA relocalizes to outside the nucleoli in budding yeast [62], and to nucleolar caps in human cells [63, 64], revealing the importance of nuclear dynamics for rDNA repair. However, the mechanisms required for this movement remain poorly understood. While more studies are required to understand the molecular mechanisms driving these dynamics, our data are consistent with a role for motor components and actin nucleation in the dynamics and repair of nucleolar DNA in *Drosophila*.

Additionally, previous studies mostly focused on the spatio-temporal regulation of heterochromatin repair in response to IR-induced DSBs. Here, our data suggest the importance of relocalization pathway components also in repair of replication-induced damage, although the nature of the damage and signaling pathways responsible for activating relocalization remain unclear. Given that fork collapse in heterochromatin is likely to induce DSBs [41], the signal triggering relocalization might rely on resection and ATR-dependent checkpoint activation, similar to DSB-induced relocalization pathways in *Drosophila* cultured cells [13].

Genome instability and mutagenesis associated with under-replicated regions of polytene chromosomes have been proposed as a contributing factor to the adaptability of *Diptera* to their environment [65]. Thus, understanding the molecular mechanisms of DSB repair in underreplicated regions can provide important insights into phenomena responsible for adaptation and evolution. These studies also pose the foundation for understanding the mechanisms of genome stability in other polytene chromosomes, including in several tissues of insects, plants, mammals, and ciliates [66, 67]. Polytene-like chromosomes also form in response to certain chemotherapy treatments [67]. Those pathological structures typically result from a single additional S-phase without separation of the replication products (polytene diplochromosomes) and can contribute to cancer formation by inducing polyploidy [67]. Thus, understanding the mechanisms of repair and stability of repeated sequences in polytene chromosomes is expected to shed light on tumorigenic processes, including cancer relapses in response to therapy.

## Statement of Author Contributions

LD performed experiments; LD, GD and IC analyzed experiments; DG and IC wrote the manuscript; IC coordinated the project.

## Acknowledgments

We would like to thank the Chiolo Lab for helpful discussions; G. Karen, J. Mason, BDSC (P40OD018537) and VDRC for fly stocks, and S. Keagy for insightful comments on the manuscript.

## Supplementary Figures

**Supplementary Figure 1:**
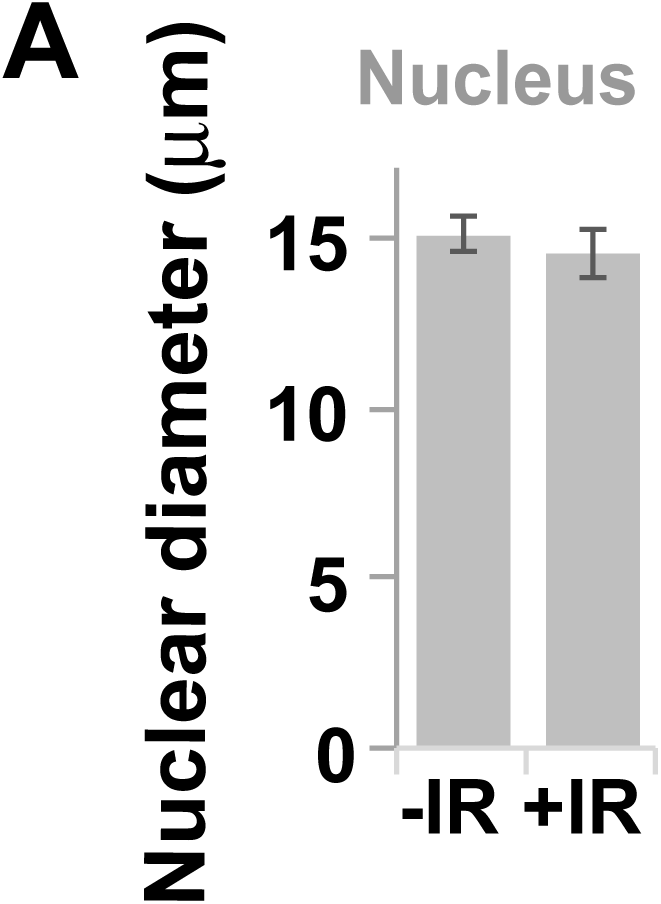
A) Quantification of the average size of the nucleus for the cells quantified in Fig. 2B.

**Supplementary Figure 2:**
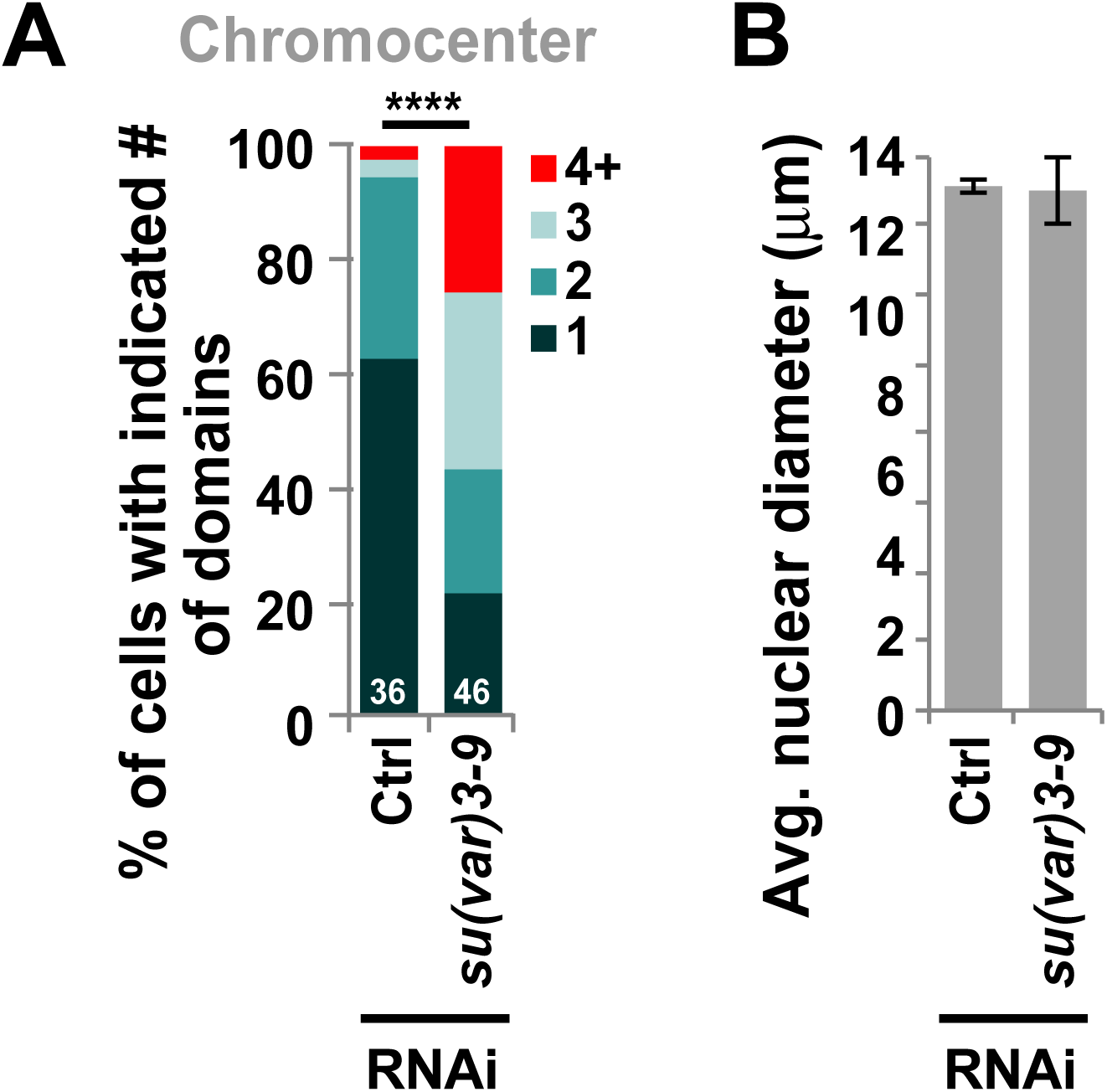
A) Quantification of the number of heterochromatin domains (H3K9me3-positive signals) in salivary gland cells of Ctrl (*yry*) or *su(var)3-9*^*null*^ mutant flies, as indicated. Numbers next to each graph indicate the number of domains in each category; numbers inside the graph are cells quantified. \*\*\**P*<0.0001, two-tailed Mann-Whitney test.

